## Supplementary material for "Drak is a potential binding partner of *Drosophila* Filamin"

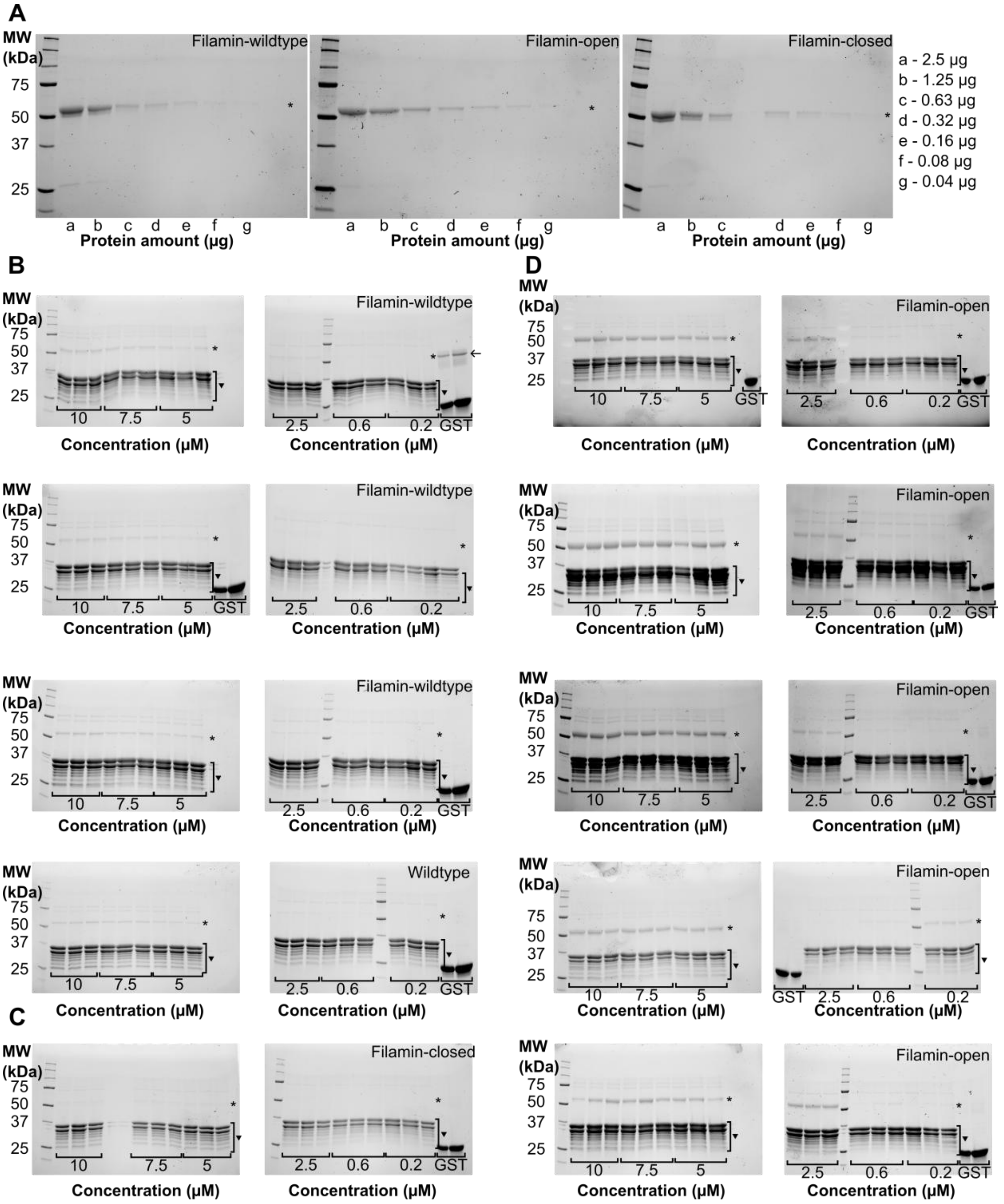


**Figure S1.** SDS-PAGE analysis of in vitro GST pull-down assay of GST-tagged C-terminal fragment of Drak kinase (▼) and five domain Drosophila Filamin fragment (*). A) A representative image of SDS-PAGE gel with varying concentrations of the purified Drosophila filamin fragments. The binding assay shows that Drak kinase interact with B) Filamin-wild type, C) Filamin-closed mutant and D) Filamin-open mutant with different affinity. The concentration of Filamin fragments used in the assay are 10 ,7.5, 5, 2.5, 0.6, and 0.2 µM. In the first right image of (B) panel, the last 2 lanes with GST (control) also have Filamin band as GST was accidentally eluted in the tube that contained leftover of Filamin used in the assay.

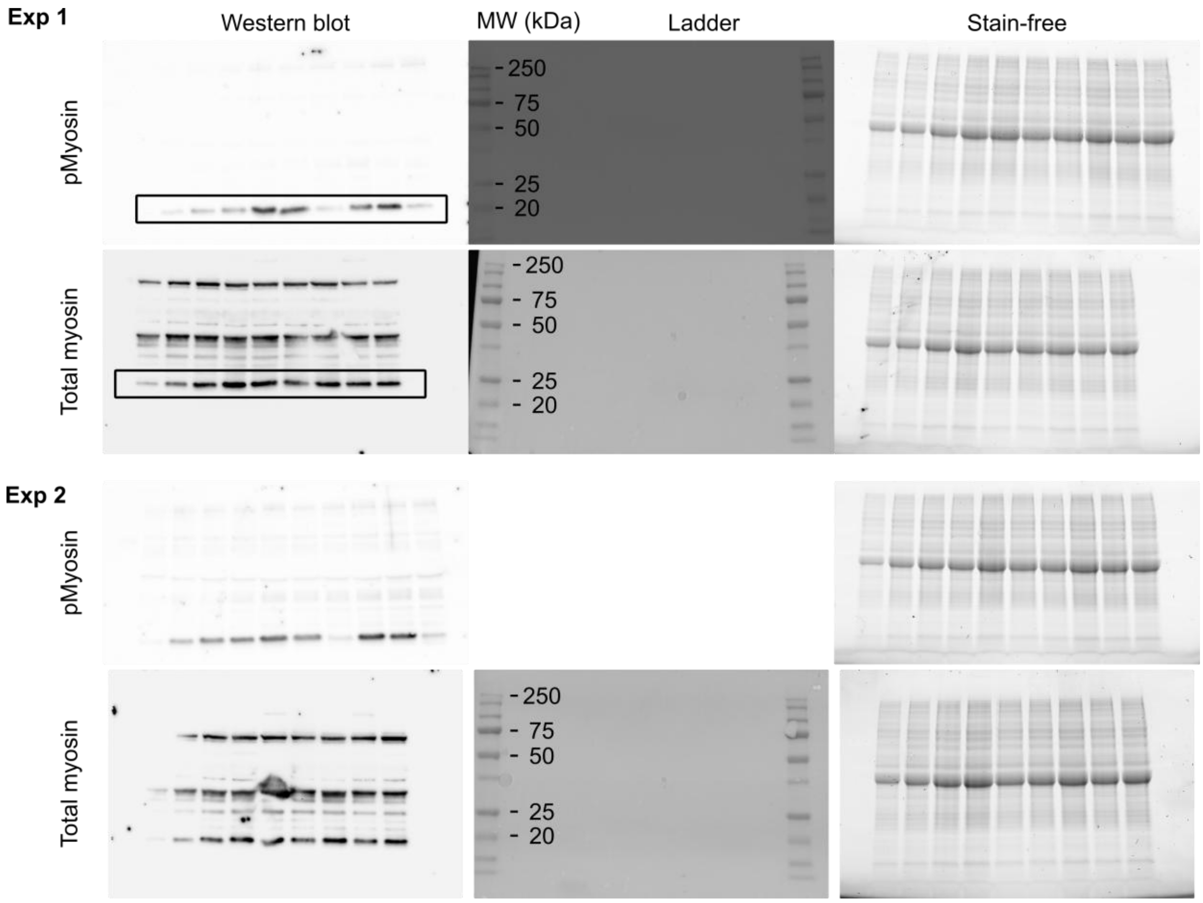


**Figure S2.** For western blot transparency. The gels and blots are fully shown in two independent experiments. Left column are western blots, middle column is ladder, right column shows the total protein amount, visualized by the stain-free fluorescence signal in the gel before transfer. The western blot bands shown in Fig. 3 are highlighted. The ladder for pMyosin is missing. Lanes for experiments 1 and 2 pMyosin: 1) Ladder, follow by WT with 2) 15, 3) 20, 4) 30, 5) 40, and 6) 50 embryos. Lanes 7) Filamin-closed, 8) Drak-KO, 9) WT 50, 10) Filamin-closed, and 11) Drak-KO had 50 embryos each. Lanes for total myosin had: 1) Ladder follow by WT with 2) 15, 3) 20, 4) 30, 5) 40 embryos. Lanes 6) Filamin-closed, 7) Drak-KO, 8) WT 50, 9) Filamin-closed, and 10) Drak-KO had 40 embryos each.

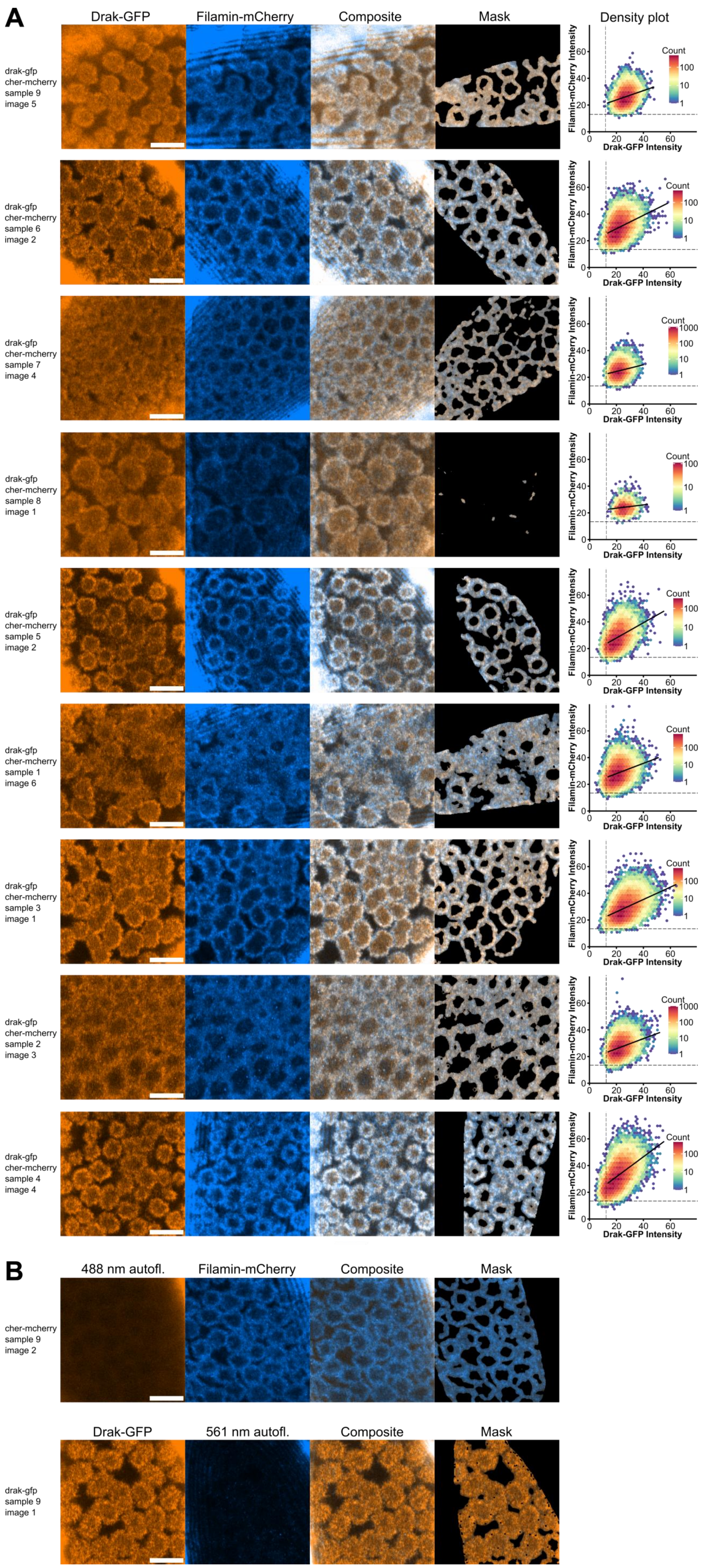
A
Drak-GFP
Filamin-mCherry
Composite
Mask
Density plot
drak-gfp
cher-mcherry
sample 9
image 5
drak-gfp
cher-mcherry
sample 6
image 2
drak-gfp
cher-mcherry
sample 7
image 4
drak-gfp
cher-mcherry
sample 8
image 1
drak-gfp
cher-mcherry
sample 5
image 2
drak-gfp
cher-mcherry
sample 1
image 6
drak-gfp
cher-mcherry
sample 3
image 1
drak-gfp
cher-mcherry
sample 2
image 3
drak-gfp
cher-mcherry
sample 4
image 4
Count
1000
100
10
1
Filamin-mCherry Intensity
Drak-GFP Intensity
0
20
40
60
B
488 nm autofl.
Filamin-mCherry
Composite
Mask
cher-mcherry
sample 9
image 2
Drak-GFP
561 nm autofl.
Composite
Mask
drak-gfp
sample 9
image 1

**Figure S3.** Representative images of all samples used for colocalization analysis at the priming phase of *Drosophila* embryo cellularization. The number of images per sample varied based on available imaging area. A total of 37 images (1–6 images/sample) were taken from 9 embryo samples. A) One representative image of each sample is shown in separate channels (Drak-GFP is orange and cher/Filamin-mCherry is blue) and as a composite. In the column labelled "Mask" the areas excluded from the ROI are shown in black. The density plot shows the intensity correlation of all pixels. Dashed vertical line shows threshold for Drak-GFP channel and horizontal line for Filamin-mCherry. The linear regression line is plotted using only pixel values that were above the threshold in both channels. B) Control images of single-fluorophore embryos that were used to determine thresholds levels for calculating colocalization parameters M1, M2, and tPCC. Masked areas were excluded as above. Upper image shown has cher/Filamin-mCherry fluorophore only used to determine threshold for Drak-GFP channel. Lower image has Drak-GFP fluorophore only to determine threshold for cher/Filamin-mCherry.

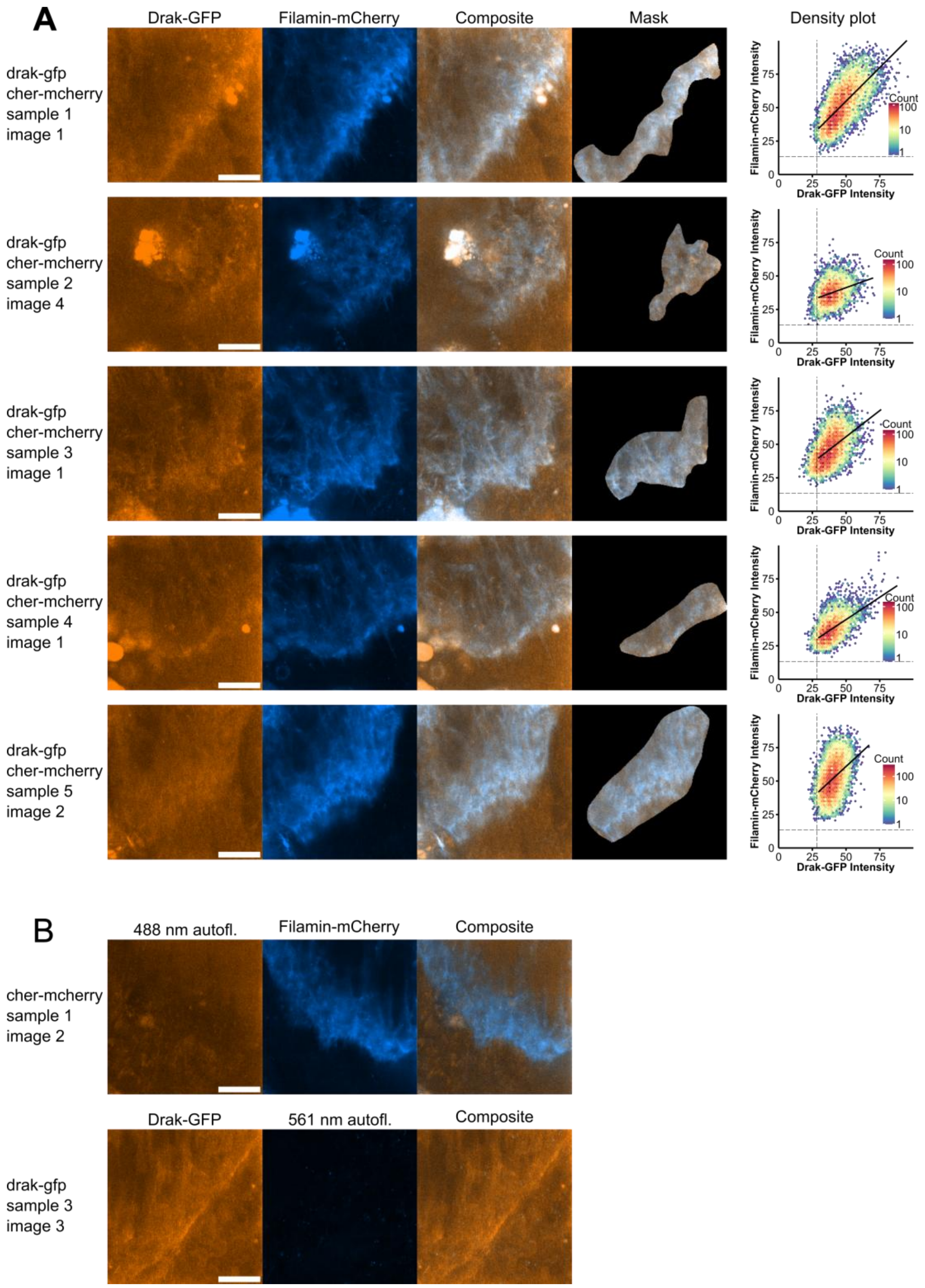


**Figure S4.** Representative images of all samples used for colocalization analysis during the tendon-myotube attachment site maturation. The number of images per sample varied based on available imaging area. A total of 17 images (1-5 images/sample) were taken from 5 pupas. A) One representative image of each sample is shown in separate channels (Drak-GFP is orange and cher/Filamin-mCherry is blue) and a composite. In the column labelled "Mask" the areas excluded from the ROI are shown in black. Density plot shows the intensity correlation of all pixels. Dashed vertical line shows threshold for Drak-GFP channel and horizontal line for Filamin-mCherry. The linear regression line is plotted using only pixel values that were above the threshold in both channels. B) Control images of single-fluorophore embryos that were used to determine thresholds levels for calculating colocalization parameters M1, M2, and tPCC. Upper image shown has cher/Filamin-mCherry fluorophore only used to determine threshold for Drak-GFP channel. Lower image has Drak-GFP fluorophore only to determine threshold for cher/Filamin-mCherry.

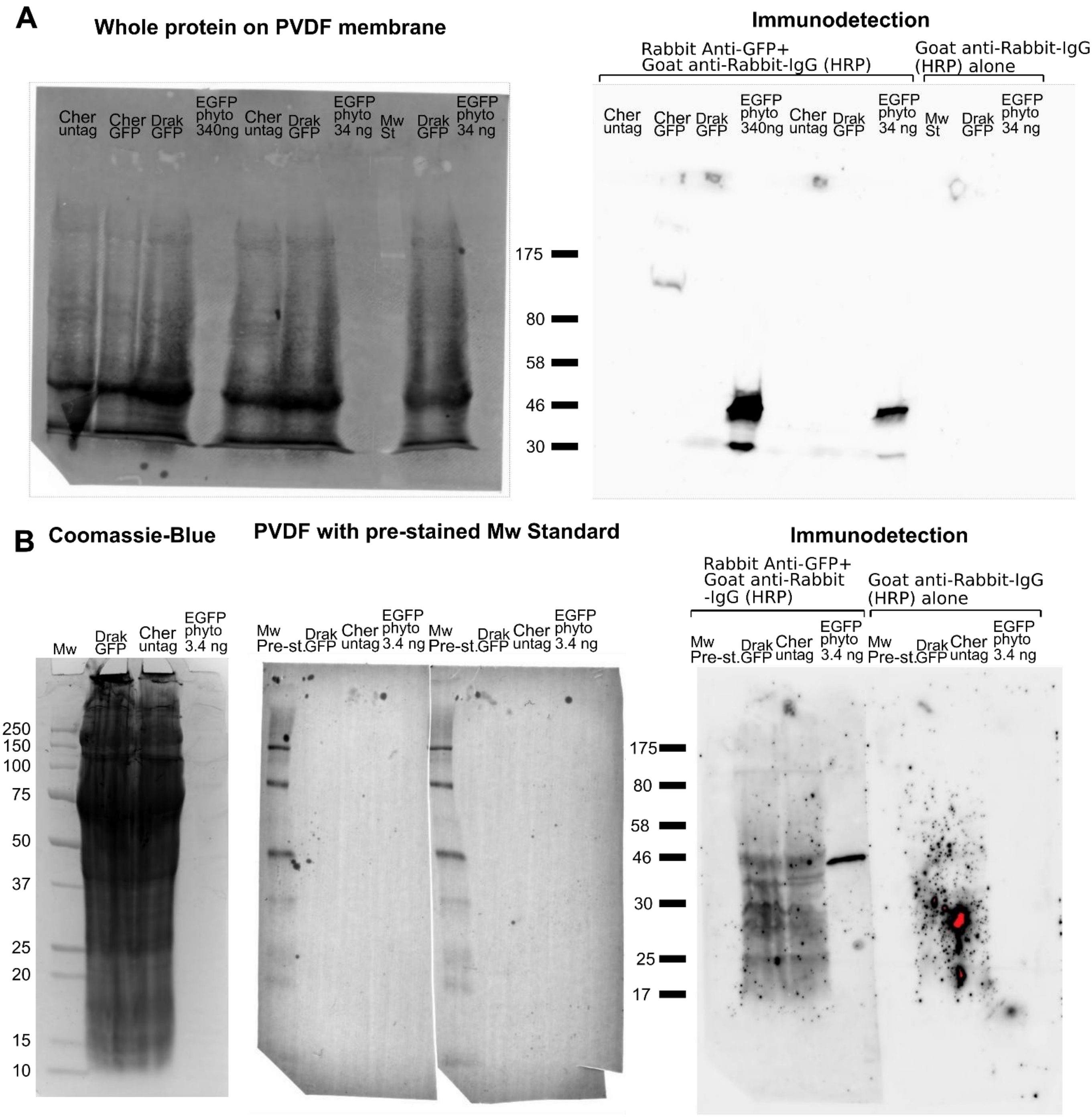


**Figure S5. Detection of GFP fusion proteins with immunoblotting**. A) Whole fly proteins separated on a 7,5 % SDS-PAGE gel. The left side shows the image of the PVDF membrane after protein transfer with the stain-free technology. The right side shows immunodetected either with anti-GFP (ab290, Abcam) and the HRP-coupled secondary antibody or the secondary antibody alone. No Drak-GFP was detected. As positive controls +test if GFP only or Drak-GFP degradation products can be detected, overloaded protein preparation of 25-28 h pupas were separated on a 12 % SDS-PAGE gel. B) The left panel shows part of the gel stained with Coomassie brilliant Blue. The same amounts of proteins were pipetted on the rest of the gel. The middle panel shows the PVDF membrane with pre-stained protein molecular weight markers and the right panel shows the same PVDF membrane immunodetected with either anti-GFP and the HRP-coupled secondary antibody or the secondary antibody alone. In both Drak-GFP pupa and non-GFP expressing pupa the same unspecific pattern is seen with anti-GFP. In both figures the sizes of the molecular weight markers are indicated as kDa.

**Table S1**. The yeast two-hybrid analysis results related to Fig 2 are given as a separate exel-file.

**Table S2.** Co-localization analysis results at the priming phase of cellularization related to Fig 4. Thresholded Manders coefficients (M1 for Drak and M2 for Filamin) and the thresholded Pearson correlation coefficient (tPCC) were calculated for each image. For validation, tPCC was also calculated for within-ROI shuffled Drak channel. Means and standard deviations were weighted by the number of informative pixels in each image: M1 by the count of Drak pixels above threshold, M2 by the count of above-threshold Filamin pixels, and tPCC by the count of pixels that exceeded the threshold in both channels (overlapping pixels).

| **Sample** | **Image** | **Overlapping Pixels** | **M1** | **M2** | **tPCC** | **Shuffled Drak tPCC** |
|---|---|---|---|---|---|---|
| 1 | 1 | 24905 | 0.992 | 0.982 | 0.271 | 0.002 |
| 1 | 2 | 19345 | 0.994 | 0.986 | 0.197 | -0.015 |
| 1 | 4 | 23685 | 0.997 | 0.973 | 0.222 | -0.007 |
| 1 | 6 | 30616 | 0.997 | 0.938 | 0.290 | -0.009 |
| 2 | 2 | 38611 | 0.997 | 0.985 | 0.322 | 0.000 |
| 2 | 3 | 41371 | 0.997 | 0.987 | 0.322 | 0.008 |
| 2 | 4 | 39766 | 0.997 | 0.983 | 0.334 | -0.007 |
| 2 | 5 | 35395 | 0.997 | 0.982 | 0.297 | 0.000 |
| 2 | 6 | 16360 | 0.998 | 0.987 | 0.254 | 0.005 |
| 3 | 1 | 33624 | 0.998 | 0.989 | 0.420 | 0.002 |
| 4 | 1 | 12338 | 0.996 | 0.873 | 0.431 | -0.008 |
| 4 | 2 | 19884 | 0.997 | 0.915 | 0.477 | 0.007 |
| 4 | 3 | 28267 | 0.996 | 0.864 | 0.388 | -0.002 |
| 4 | 4 | 30309 | 0.998 | 0.902 | 0.493 | -0.005 |
| 4 | 5 | 32522 | 0.999 | 0.944 | 0.517 | -0.006 |
| 4 | 6 | 28942 | 0.998 | 0.951 | 0.484 | -0.002 |
| 4 | 7 | 28338 | 0.998 | 0.958 | 0.429 | -0.001 |
| 4 | 8 | 23927 | 0.996 | 0.880 | 0.319 | 0.006 |
| 5 | 1 | 5775 | 0.994 | 0.894 | 0.345 | 0.000 |
| 5 | 2 | 17325 | 0.996 | 0.929 | 0.396 | 0.015 |
| 5 | 3 | 10778 | 0.998 | 0.930 | 0.521 | -0.002 |
| 6 | 1 | 12017 | 0.998 | 0.955 | 0.371 | 0.007 |
| 6 | 2 | 27819 | 0.999 | 0.971 | 0.368 | 0.001 |
| 6 | 3 | 15845 | 0.998 | 0.969 | 0.368 | 0.008 |
| 7 | 1 | 7373 | 0.999 | 0.994 | 0.255 | -0.003 |
| 7 | 2 | 6303 | 0.999 | 0.979 | 0.115 | -0.003 |
| 7 | 3 | 17344 | 0.999 | 0.981 | 0.206 | -0.006 |
| 7 | 4 | 26302 | 0.998 | 0.994 | 0.218 | 0.009 |
| 8 | 1 | 2869 | 0.999 | 1.000 | 0.122 | -0.006 |
| 8 | 2 | 388 | 1.000 | 1.000 | 0.110 | 0.012 |
| 8 | 3 | 1249 | 0.999 | 1.000 | 0.220 | 0.036 |
| 8 | 4 | 1292 | 0.999 | 1.000 | 0.155 | -0.013 |
| 9 | 1 | 4591 | 0.998 | 0.999 | 0.241 | 0.002 |
| 9 | 2 | 4491 | 0.997 | 1.000 | 0.255 | 0.002 |
| 9 | 3 | 13140 | 0.998 | 1.000 | 0.326 | -0.001 |
| 9 | 4 | 10714 | 0.998 | 1.000 | 0.377 | 0.004 |
| 9 | 5 | 19034 | 0.999 | 1.000 | 0.284 | -0.010 |
| | | **mean** | 0.998 | 0.970 | 0.325 | |
| | | **p-value** (one-sample t-test) | 2.2e-16 | 2.26e-13 | 2.33e-5 | |

**Table S3.** Co-localization analysis at tendons during tendon-myotube attachment site maturation in developing indirect flight muscles, related to Fig 7. For each image we calculated the thresholded Manders coefficients (M1 for Drak and M2 for Filamin) and the thresholded Pearson correlation coefficient (tPCC). For validation, tPCC was also calculated for within-ROI shuffled Drak channel. Means and standard deviations were weighted by the number of informative pixels in each image: M1 by the count of Drak pixels above threshold, M2 by the count of above-threshold Filamin pixels, and tPCC by the count of pixels that exceeded the threshold in both channels (overlapping pixels).

| Sample | Image | Overlappin Pixels | M1 | M2 | tPCC | Shuffled Drak tPCC |
|---|---|---|---|---|---|---|
| 1 | 1 | 14442 | 1.000 | 0.997 | 0.673 | -0.010 |
| 1 | 2 | 13541 | 1.000 | 0.999 | 0.593 | -0.009 |
| 2 | 1 | 6455 | 1.000 | 0.963 | 0.289 | 0.012 |
| 2 | 2 | 5259 | 1.000 | 0.763 | 0.311 | -0.027 |
| 2 | 3 | 4248 | 1.000 | 0.943 | 0.303 | -0.007 |
| 2 | 4 | 8187 | 1.000 | 0.967 | 0.303 | -0.001 |
| 2 | 5 | 6927 | 1.000 | 0.909 | 0.512 | 0.006 |
| 3 | 1 | 13429 | 1.000 | 0.915 | 0.473 | -0.019 |
| 3 | 2 | 7787 | 1.000 | 0.972 | 0.373 | 0.015 |
| 4 | 1 | 8486 | 1.000 | 0.963 | 0.562 | 0.009 |
| 4 | 2 | 3562 | 0.999 | 0.422 | 0.318 | -0.023 |
| 4 | 3 | 6936 | 1.000 | 0.429 | 0.152 | -0.015 |
| 4 | 4 | 6146 | 1.000 | 0.582 | 0.272 | -0.022 |
| 5 | 1 | 19778 | 1.000 | 0.984 | 0.431 | 0.002 |
| 5 | 2 | 22319 | 1.000 | 0.989 | 0.419 | 0.001 |
| 5 | 3 | 12866 | 1.000 | 0.896 | 0.462 | 0.001 |
| 5 | 4 | 10871 | 0.997 | 0.849 | 0.522 | 0.006 |
| | | **mean** | 0.999 | 0.870 | 0.450 | |
| | | **p-value** (one-sample t-test) | 4.97e-16 | 0.00035 | 0.00241 | |

**Movie S1.** This movie is related to Fig. 6. The compaction and elongation phase of developing of IFMs in *Drosophila*, visualized with Drak-GFP (orange) and Filamin-mCherry (blue). The video shows a single confocal section with the 60x objective. The time is after pupal formation with time interval of 1 min and video speed of 30 min/s.
